# Measuring hospital spatial wingspan by using a discrete choice model with utility-threshold

**DOI:** 10.1101/468645

**Authors:** Saley Issa, Ribatet Mathieu, Molinari Nicolas

## Abstract

Policy makers increasingly rely on hospital competition to incentivize patients to choose high-value care. Travel distance is one of the most important drivers of patients’ decision. The paper presents a method to numerically measure, for a given hospital, the distance beyond which no patient is expected to choose the hospital for treatment by using a new approach in discrete choice models. To illustrate, we compared 3 hospitals attractiveness related to this distance for asthma patients admissions in 2009 in Hérault (France), showing, as expected, CHU Montpellier is the one with the most important spatial wingspan. For estimation, Monte Carlo Markov Chain (MCMC) methods are used.

## Introduction

It is often advocated that competition between hospitals may improve efficiency and quality of health care [1]. In fact, in theory, due to increasing competition, hospitals decrease production costs and improve the quality of health care delivered so as to attract new patients. Factors of hospital choice include price, quality, distance or travel time (easy access), waiting time, provider network and others. Their relative importance differs according to the market characteristics and the regulatory context [2]. Early studies identified distance or travel time as the major factor negatively affecting hospital choice [3], [4] even if sensitivity to distance varies with patient characteristics (age, ethnicity, income and religion), admission types (i.e., stronger effect of distance for common procedures) and hospital type (i.e., weaker effect for admissions to larger specialized hospitals) [5].

As in many European countries, French hospitals are financed through Diagnosis Related Groups based on a prospective payment system. Regarding acute care, this scheme has been fully implemented in 2005 for private for-profit hospital budgets and in 2008 for public hospital budgets. The reform was intended to improve efficiency and fairness in financing and also to increase competition between and within the public and private sectors [6]. Information about performance and quality of hospitals is not yet published in France, and waiting times are currently not identified as an issue. Actually, hospital choice is made by patients themselves, following advice from their general practitioners. Since general practitioners do not face any financial incentives to refer their patients to a given hospital, we can assume that they take into account patients’ preferences among other factors. Thus, we can hypothesize that reputation [7], as perceived by the patients and the general practitioners is partially reflecting the quality of care and easy access. In these circumstances, travel distance certainly plays a key role in patients’ decision [8]. Further, in areas where there are many hospitals within short distance, the attractiveness of a hospital (easy access, care quality, …) can be measured by the distance travelled by patients to get there. That is to say, longer the distance travelled, the more attractive the hospital.

In this context, a spatial modeling on hospitals choice may be very useful to propose a classification of hospitals (most to least attractive). From a policy perspective, these findings have implications for regulation. For instance, the effect of making the quality of care information publicly available may not have the expected impact in areas where facilities are scarce and distant from one to another. More generally, the predominant effect of distance on hospitals choice should be acknowledged and taken into consideration before implementing any reforms aimed at incentivizing hospitals competitiveness.

In this work, we present a method to numerically measure the distance beyond which no patient is expected to choose the hospital for treatment by using a new approach in discrete choice models. This distance is used to compare hospitals attractiveness.

For illustration, we used patients admissions for asthma in 3 hospitals of Hérault that have been registered in 2009.

The paper is organized as follows. Section **Data** describes the data we used for illustration while Section **Proposed Approach** explains the approach we introduced to cope with the issue we encountered with the RUMs. An application of the approach is presented in Section **Illustration** and the paper ends with a brief discussion.

## Data

The data we used in this work, were obtained in a fully anonymized and de-identified manner from the *Programme de Médicalisation des Systèmes d’lnformation* (PMSI). The PMSI is an administrative dataset recording all inpatient admissions in French private and public hospitals covering all social health insurance schemes. It is intended to describe the medical activities of hospitals. Information about the patient and the hospital of stay such as the patient desease, his place of residence, location of the hospital, type of hospital, are held in the dataset.

For our case study, we used data about asthma hospital patients stays in Languedoc–Roussillon, a region known to have a high competitive health care market in France. Located in the South of France, (Fig 1), the region consists of five departments: Aude, Gard, Hérault, Lozère and Pyrénées-Orientales. It extends from 42° to 44.5° Latitude North and between 1.5° to 5° Longitude East. The Languedoc–Roussillon region has a population of about 3 million people, which includes both rural and metropolitan areas. It is the second region with the highest population growth (1.1%) in France (about 0.4%) due essentially to immigrants and more than 16% of the population are retirees.

**Fig 1.**
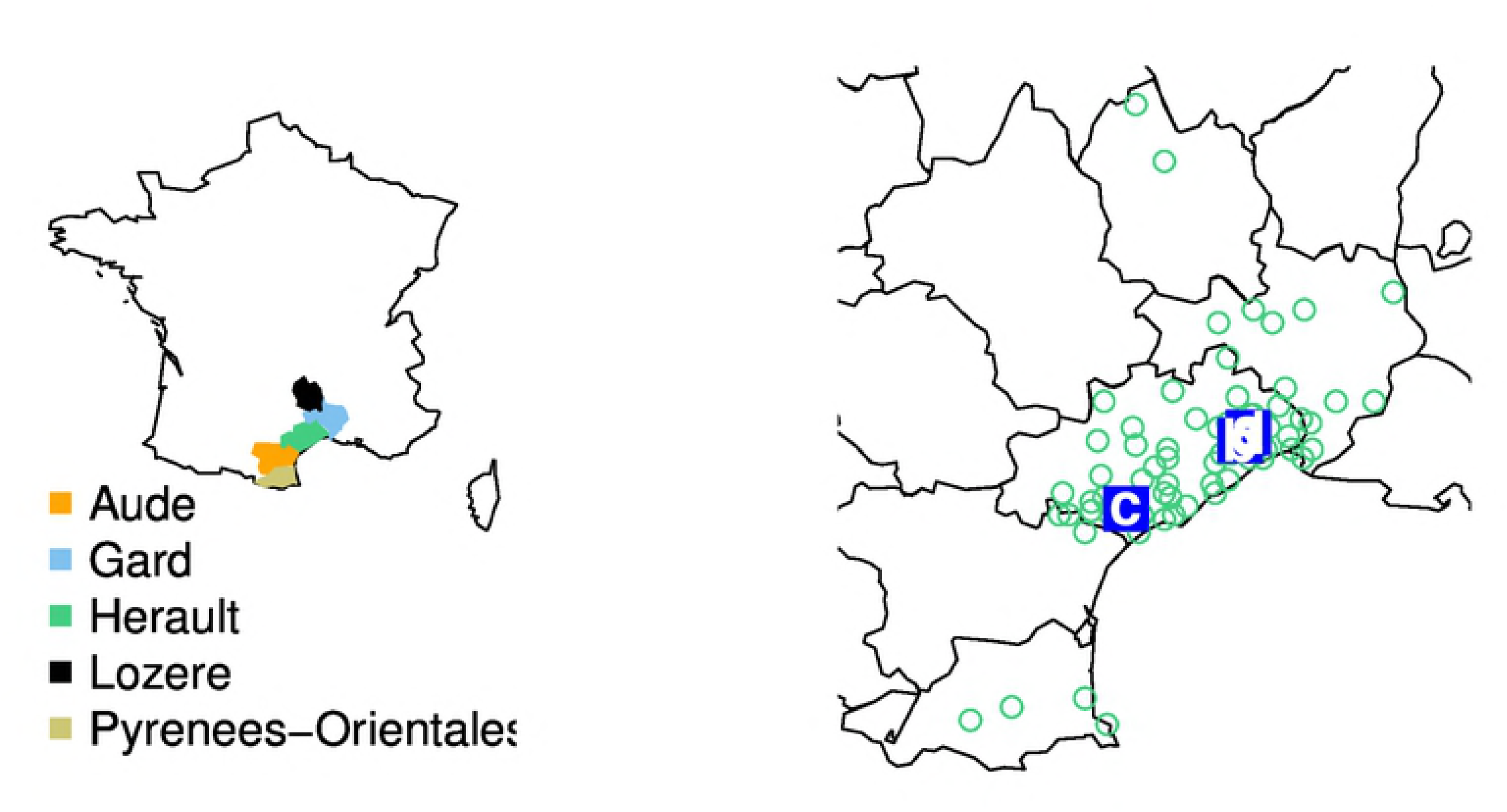
Mapping of the illustration data. The left map shows where the Region Languedoc-Roussillon is located in France. The picture on the right displays locations of the hospitals (a: CHU Montpellier, b: Clinique le Millénaire and c: CH Béziers) and localities (green points) where at least one resident admission is reported in these hospitals

In 2009, 1,289 hospital patient stays have been recorded for asthma in this region as reported in [9]. The study focused on admissions in 3 hospitals of Hérault: one is private for-profit and 2 are not-for-profit. Table 1 displays the hospitals with the number of stays.

**Table 1.** Number of asthma stays registrated per each of the 3 hospitals in 2009

| Hospital | number of asthma patient stays |
| --- | --- |
| CH Béziers | 79 |
| CHU Montpellier | 127 |
| Clinique Le Millénaire | 23 |

For each of these 3 hospitals, we extracted from the dataset their geographic coordinates, the number of patients admitted per locality and the geographic coordinates of these localities. From the geographic coordinates, we computed euclidean distance between the hospital and patient locations, used as travel distance, because having road distance between localities can be tricky.

## Proposed approach

### Motivation

Patients admissions to hospitals can be seen as a situation of choice where agents (here patients) have to choose among a set of products or alternatives (here hospitals for a given therapy); therefore discrete choice models can be used to model the data. Since it seems reasonable to consider that a patient chooses an hospital when obtaining a certain utility of doing so, Random Utility models (RUMs) [10] can be used.

The concept of these models is that for any given alternative *j* from *J* alternatives in competition, an agent *n* is expected to obtain a certain utility *U_nj_* of choosing the alternative. This utility *U_nj_* is often decomposed in the sum of 2 parts:

a. *V_nj_* (called representative utility), the part of utility described by the observed variables influencing the choice of the agent;
b. *ϵ_nj_*, the part of utility related to unobserved variables also influencing the choice of the agent,

i.e, *U_nj_* = *V_nj_* + *ϵ_nj_*. By considering *ϵ_nj_*, *j* = 1, …, *J*, random, thus assigning them a distribution *f* and under the assumption that the agent chooses the alternative with the greatest utility, the probability of choosing an alternative *j* by the agent *n* can be derived as

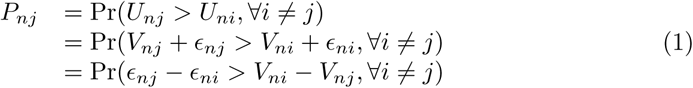

According to the specification of *f* (.), different specific models are obtained. For example, the Logit model [11] is obtained under the assumption that *ϵ_nj_* are independent copies of a Gumbel distribution, i.e,

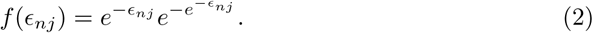

This model is by far the most widely used among RUMs due to some of its properties. But through the model use, limitations arise: independence of unknown parts may not always hold; Logit can not handle some cases of taste variation or some panel data. Various models have been developed to cope with these limitations. We can cite the Generalized Extreme Value distribution among which the most well-known is the Nested-Logit model [12]. It is derived under the assumption that the cumulative joint distribution of *ϵ_nj_*, *j* = 1, …, *J* is

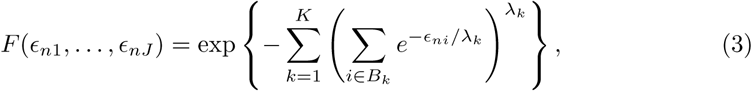

where *B_k_* is a subset of the *J* alternatives partitioned in *K* non-overlapping subsets called Nests so as for 2 alternatives *j* and *m* in a same Nest *B_k_*, *ϵ_nj_* and *ϵ_nm_* are correlated and for 2 alternatives *j* and *m* in 2 distinct Nests, respectively *B_k_* and *B_l_*, *k* ≠ l, *cov*(*ϵ_nj_*, *ϵ_nm_*) = 0. The parameter *λ_k_* measures the degree of independence of the *ϵ_nj_*-in Nest *B_k_*; the lower the value of *λ_k_*, the more correlated the *ϵ_nj_*-and the higher its value, the less correlated the unobserved factors. The value of *λ_k_* = 1 corresponds to Logit model.

Probit ([13] for binary case) is developped under the assumption that

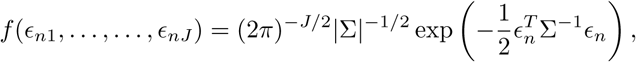

with *ϵ_n_* = (*ϵ*_*nJ*1_,…, *ϵ_nJ_*)^T^ and Σ the covariance matrix. The problem with this model is that the probability of choice is not closed-form and hence, requires simulations.

Mixed Logit [14] and [15], more flexible [16], is so as if *β* is the unknown parameter of *V_nj_*, then the choice probability is

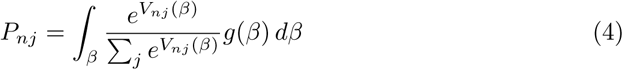

where *g* can be any density function on *β*.

In this study, we have patients admissions for asthma in 3 hospitals of Hérault that have been registered in 2009. To use RUMs, we define the utility that may obtain a patient *n* from choosing hospital *j* as

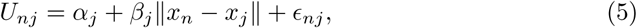

that is to say

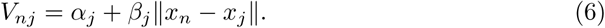

*α_j_* and *β_j_* are unknown parameters specific to the hospital *j* and ∥*x_n_* – *x_j_*∥, the euclidean distance between patient *n* and hospital *j* locations. *β_j_* controls how *V_nj_* varies according to the distance. By assessing *α_j_* positive and utility decreasing (thus *β_j_* negative) by far one moves away the hospital *j*, the ratio

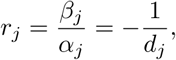

where *d_j_* is the distance beyond which *V_nj_* ≤ 0. This distance *d_j_* called wingspan, can be interpreted as the distance beyond which no patient is expected to choose the hospital considering the distance since the representative utility is negative.

However, first, *r_j_* can not be estimated with RUMs. In fact, the constants *α_j_* can not be estimated in the expression

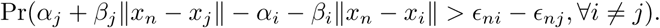

Only the difference *α_j_* – *α_i_* can be estimated; and there is an infinity of constants *μ_j_* and *α_i_* giving the same difference. One more problem in estimating of *r_j_*, is that for 2 hospitals *j* and *i* in a same locality, *β_j_* and *β_i_* can not be estimated but their difference because ∥*x_n_* – *x_j_* ∥ = ∥*x_n_* – *x_i_*∥. For instance, it is the case for CHU Montpellier and Clinique le Millénaire in our study, which are both located in Montpellier. These two issues make that *r_j_* can not be estimated with the RUMs, though knowing this ratio is needed in this study in order to compare the attractiveness of the hospitals.

Second, the assumption of utility-maximization in choosing a hospital for treatment can be not true for some patients (bounded rationality).

So, to estimate *r_j_* and cope with the situation where patients do not maximize the utility, we propose a new approach. In our approach, the utility is really considered deterministic and the assumption of utility maximization, which is criticized by many psychologists is released for a more loose assumption by introducing a utility-threshold: any alternative whose utility for the agent *n* exceeds a certain threshold *S_n_*, can be chosen by this agent.

### The approach

In order to cope with the estimation issue of ratio mentioned above, we present another approach where the assumption of utility maximization used to derive the choice probability is released. In fact, when an agent is faced with a choice among several alternatives, he is often not sure which alternative he should select, and does not always make the same choice under seemingly identical conditions, as noted by Tversky [17]. Therefore, it is strongly possible that an agent does not always maximize the utility when choosing an alternative. As an agent does not always choose the alternative with the greatest utility for him and as it is unlikely he chooses an alternative that he thinks useless or harmful for him, we can suppose that he chooses an alternative from the moment the utility he may obtain considering some aspects covers his certain need (Bounded rationality). It is this certain need that we call a threshold of utility. It is a minimum of utility that may demand the agent from an alternative. It can vary due to socio-demographic variables, the type of alternatives, and other unknown factors. Since we don’t know how exactly it varies, though that these socio-demographic variables can make an agent to be more or less demanding, we treat it random and assign it a distribution. Thus, any alternative probability choice of the agent considering some aspects can be derived.

To formalize the idea, consider that we have an agent *n* facing choice among *J* alternatives. As for random utility models, these alternatives are in finite number. They do not have to be mutually exclusive, but just exhaustive in the sense that any choice of the agent may be met by the combination of these alternatives. Let *U_nj_* be the utility that may obtain the agent *n* considering some aspects from choosing the alternative *j*. The agent can choose the alternative if only the utility he may obtain from doing so, exceeds a minimum of utility (called utility-threshold) *S_n_*, i.e, *U_nj_* > *S_n_*. The *S_n_* is treated as random and assigned a probability density function *f*. The probability of an alternative *j* to be chosen by the agent *n* is defined as just

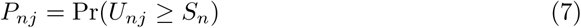

The greater *U_nj_* is, compared to *S_n_*, the more likely the alternative is going to be chosen. Therefore, the alternative with the greatest utility considering the aspects has the most chance to be chosen.

Considering that it is more realistic for an agent to choose an alternative if having a benefit, the threshold is expected to be positive. This utility-threshold can be supposed as a conjonction of many independant factors (incomes, religion, education…). Therefore, we can assess that the logarithm of *S_n_* has a normal distribution with parameters (*μ_n_*, *σ_n_*), i.e,

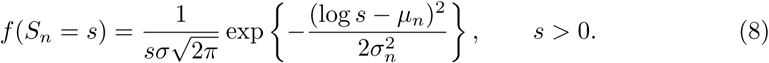

Thus,

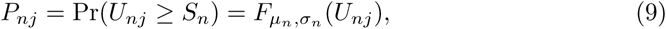

where *F*_*μn*,*σn*_ is the cumulative log-normal distribution with parameters (*μ_n_*,*σ_n_*). But in general, it is known that absolute utility can not be measured, so *μ* may not be known. We know that from the constraint, exp(*μ_n_*) is from zero to infinity. So we divide *U_n_* by exp(*μ_n_*) and re-state the probability choice so as

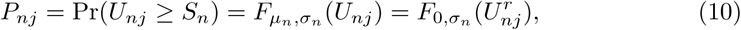

where 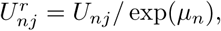 a relative utility which is to be estimated. For Eq (10), in fact, it is easy to show that for any *x*

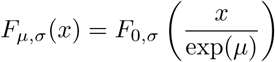

for a log-normal distribution.

The probability of an agent choice can be written as

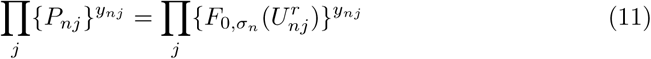

where *y_nj_*=1 if *j* is chosen by *n* and 0 otherwise.

For a sample of *N* agents choosing among *J* alternatives and considering that the *S_n_* are independent, the likelihood for *A*, the set of observed choices, is

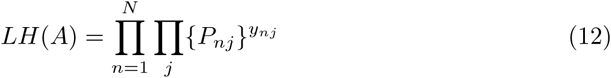

For unkown parameters estimation, likelihood maximization can be used; but in this paper, a Bayesian framework is adopted.

## Illustration

Recall that the aim of this study is to compare the attractiveness of the hospitals according to the travel distance made by patients. In fact, we have the number of patients by locality, hospitals where they went and eucludean distances obtained from geographic coordinates of the localities.

### Model under the proposed approach

Consider that we have *N* patients who face choice among *J* hospitals. We suppose that the only observed variable that may determine their choice is travel distance. Therefore, we define the utility that may obtain the patient *n* considering the distance from choosing hospital *j* as

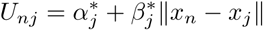

where 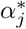 is positive, *x_n_*, *x_j_*, respectively geographic coordinates of the patient *n* and hospital *j* locations and ∥*x_n_* – *x_j_*∥, the eucludean distance between these locations. The parameter 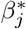 exhibits the effect of distance. Since the utility of a hospital is supposed to reduce as one moves away, the parameter 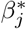 is expected to be negative. From the assumption that the threshold *S_n_* is distributed log-normal, then for *U_nj_* ≤ 0, the probability that the patient n chooses the hospital j is zero. It means that beyond the distance *d_j_* so as 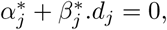 no patient is supposed to choose the hospital *j* for treatment. This distance will be used to compare hospitals attractiveness. The bigger this distance is, the more attractive the hospital is. By estimating the ratio

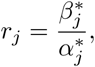

we can subsequently estimate *d_j_*.

The main target is now to estimate the ratio *r_j_* for the 3 hospitals with 229 patients stays (Table 1) by using the approach above in a Bayesian framework.

Denote *A*, the set of choices actually observed. To reduce over parameterization, we assume that the parameters of the log-normal distribution are the same for all the patients since people in a same community share almost the same characteristics, i.e, (*μ_n_*,*σ_n_*) = (*μ*, *σ*), ∀*n*. Then we define

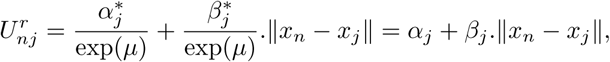

the relative utility, without impact on *r_j_* because

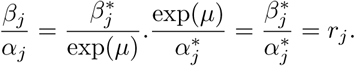

Supposing that *S_n_*, *n* = 1, …, 229 are independent, then the probability to have *A*, the set of the observed choices, when, *σ*, *α_j_*, *β_j_*, *j* = 1, 2, 3 are known can be expressed as

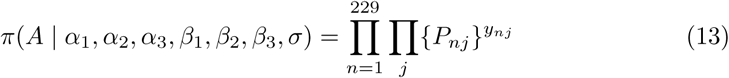

where *P_nj_* = *F*_0,*σ*_(*α_j_* + *β_j_*·∥*x_n_* – *x_j_*∥) and *F*_0,*σ*_, cumulative log-normal distribution with parameters (0, *σ*). *y_nj_* = 1 when the agent *n* chooses the alternative *j* and zero otherwise.

It can be good to use joint distribution for some of the parameters to take into account the presence of other alternatives but we suppose them all independent. Hence, for each of *α_j_*, *β_j_* and *σ*, we set a prior distribution. For *α_j_*, a uniform distribution can be used. But since it is unlikely for *α_j_* to be infinite and we don’t know its exact upper bound, thus, we prefer to use an Inverse-Gamma distribution with scale and shape parameters equal to 1. Also an Inverse-Gamma is used for *σ* but the scale and shape parameters equal 2 and for *β_j_*, a Uniform *U*(–2,0), as prior distributions.

### Estimation methods

For simplicity, denote *α* = (*α*_1_,*α*_2_,*α*_3_), *β* = (*β*_1_,*β*_2_,*β*_3_), *α*_–*j*_, *α* without the component *α_j_* and *β*_–*j*_, *β* without the component *β_j_*.

To compute *r_j_*, a sample of the parameters has to be drawn from the posterior distribution *π*(*α*, *β*, *σ* | *A*).

MCMC algorithms such as Gibbs sampling [18] are often used to have approximate draws from this kind of distributions. For Gibbs sampling, parameters conditional distributions given in Eq 14 whose detailed expressions are given in Appendix are to be used. But, these distributions are not closed-form; which means that drawing from these distributions is not straightforward.

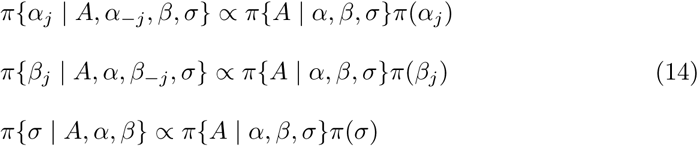

So, we used Metropolis-Hasting steps [19] within a Gibbs sampler to update each parameter. For proposal ditribution, normal distribution was used for *β_j_*, and a log-normal was used for *α_j_* and *σ*, for *j*=1,2,3.

## Results

To collect sample draws, we ran 100,000 iterations with the MCMC codes. We removed the first 20,000 iterations considered as burn-in period of the chain, then in the remaining iterations, we kept the draws by step of 10 iterations to reduce the dependence within draws. To check the convergence of the MCMC chains obtained, we made plots of these chains for each parameter and traced the corresponding conditional distribution. For Clinique le Millénaire, the chains of *α_j_* and *β_j_* are variable and not mixing well. It is likely due to the lack of enough locations of observations: in fact, there are 13 localities without zero admission. So, we produced 10 MCMC chains and though the variability of *α_j_* and *β_j_* of Clinique le Millénaire, the computed mean of *r_j_* is stable for all the 3 hospitals as reported in Table 2; which means that this ratio can be used for comparison. The table 3 gives the computed average distance at which the utility provided by the hospital is zero; which means that beyond this distance, no asthma sick is expected to come in the hospital according to the data. For recall this distance *d_j_* is obtained by equaling 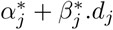 to zero, that’s to say 1 + *r_j_*·*d_j_* = 0.

**Table 2.** Table of the computed mean of *r_j_* and its Standard Error (SE) over 10 MCMC chains for the asthma dataset

| Hospital | $r_j$ mean value | SE |
| --- | --- | --- |
| CH Béziers | -0.0042 | 0.0004 |
| CHU Montpellier | -0.0024 | 0.0001 |
| Clinique le Millénaire | -0.0166 | 0.0003 |

**Table 3.** Table of average distance and its Standard Error (SE) over 10 MCMC chains. Beyond this distance no patient is expected to come in the hospital for treatment due to asthma

| Hospital | $d_j$ (km) | Std. Err. |
| --- | --- | --- |
| CH Béziers | 239 | 17 |
| CHU Montpellier | 426 | 17 |
| Clinique le Millénaire | 60.4 | 1.2 |

The results clearly show that the CHU Montpellier is the most attractive and with an important wingspan that goes beyond the Languedoc-Roussillon, the administrive zone of the center, even if this wingspan has to be relativized due to the distance used and the assumptions made on the sample. Its attractiveness is not surprising. CHU Montpellier is a well-known center providing a high level healthcare service in Languedoc Roussillon and has big accomodation capacity. It has also easy access due to transportation facilities: Highway A9, railways (TGV and TER) and many other roads. The rank of CH Béziers is plausible because it is in reality the second health center in Hérault with lower activity than CHU Montpellier.

Clinique Le Millénaire is most negatively affected by distance and with a wingspan that does not go beyond the department of Hérault. The rank of Clinique le Millénaire can be explained by the fact it is a private hospital, more specialized in acute health care like surgery and with less accommodation capacity. The results are in adequacy with what we expected for the attractiveness of the hospitals in competition.

Though the results obtained seem to be plausible, in Bayesian analysis, it is important to know how sensitive they are to the choice of the prior distributions. In our illustration, it is good to know how the *r_j_* is sensitive to the choice of the different priors; mainly the choice of the priors of *α_j_* and *β_j_*. So, to check the sensitivity, we keep one of 2 priors constant and make vary the hyperparameters of the other. For *β_j_*, we used unif(−5,0), unif(−10,0) and unif(−20,0) and no sensible variation are observed. But when we make vary the prior of *α_j_*, the mean of *α_j_* becomes smaller when its prior is more informative and bigger in the opposite. And by the way, it changes the mean of *β_j_*. Even if the change is not very big, the mean of *r_j_* seems to become bigger for CH Montpellier and smaller for Clinque le Millénaire (which has a small number of admissions), when the prior of *α_j_* is more informative as reported in Table 4. But when using less informative prior for *α_j_*, we noticed that the ratio for CHU Montpellier not stable. So, caution has to be used when choosing the prior of *α_j_*.

**Table 4.** Table of the computed mean of *r_j_* over 10 MCMC chains according to the Inverse-Gamma (*IG*) used as distribution of *α_j_*

| | $IG(2, 2)$ | $IG(3, 3)$ | $IG(5, 5)$ |
| --- | --- | --- | --- |
| CH Béziers | -0.004 | -0.004 | -0.004 |
| CHU Montpellier | -0.002 | -0.002 | -0.002 |
| Clinique le Millénaire | -0.017 | -0.017 | -0.017 |

## Discussions-Conclusions

In this paper, we presented a method to measure hospital spatial attractiveness by using a new approach in discrete choice models. This approach releases the assumption of utility-maximization which seems sometimes not realistic for a more loose assumption. The assumption supposes a threshold of utility above which an agent can choose an alternative. Application of the approach is done to asthma patients admissions in 2009 for 3 hospital of Hérault. Plausible results are obtained even if they have to be relativized to the assumptions made on the sample.

But, in spite of the obtained plausible results, application to other discrete choice data has to be led to further study of the approach and comparison with other models in order to know its power and limitations. For instance, utility correlation between alternatives (in a hierarchical model) can be considered to improve the model.

## Appendix: Conditional distributions derivation

In this section, we give the derivation of parameters conditional distributions used in the Gibbs sampler (see section **Illustration**). Then the explicit form of the distributions is obtained by replacing every term of these distributions by its mathematical expression.

For recall, Bayes formula is

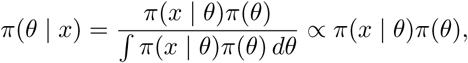

where *π*(*θ*| *x*) is called posterior distribution, *π*(*θ*) prior distribution and *π*(*x* | *θ*), the likelihood.

Consider *π*(*A*, *α*, *β*, *σ*) as the joint distribution of *A*, *α*, *β* (as defined in section **Illustration**) and *σ*.

Using the definition of conditional probabilities and assumption of independence of the parameters within their prior distributions, we show that

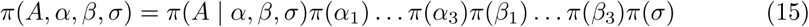

Then using Bayes formula and the Eq (15), we obtain

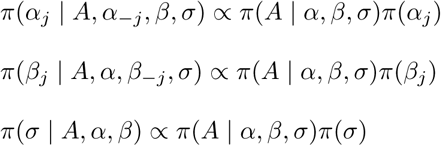

which are the conditional distributions to be used in the Gibbs sampling. To have an explicit form of these distributions, every term in the right members has to be replaced by its mathematical expressions which are:

a. for prior distributions,

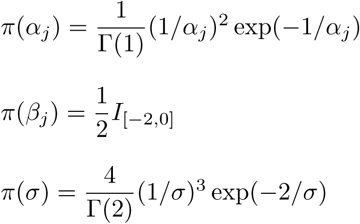
b. and for the likelihood,

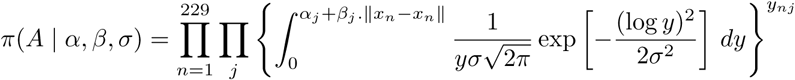

where 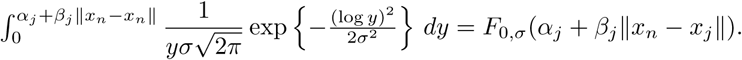 Therefore, we obtain

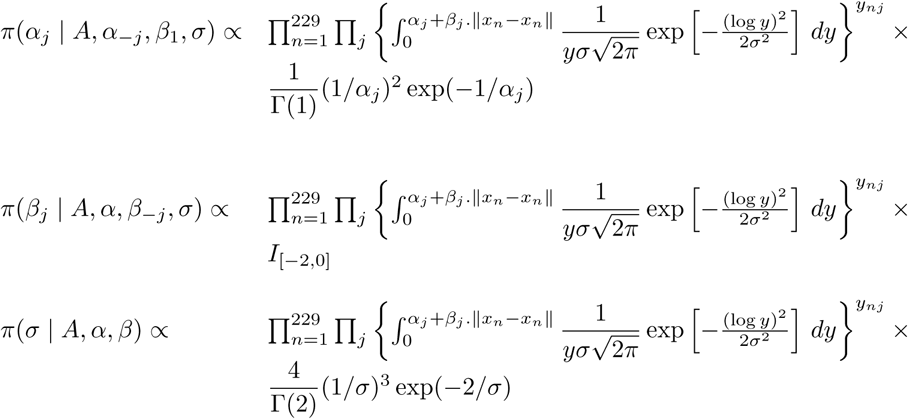

